# An optimized method to visualize lipid droplets in brain tissue demonstrates their substantial accumulation in aged brains

**DOI:** 10.1101/2024.06.12.598519

**Authors:** Francesco Petrelli, Alicia Rey, Diana Panfilova, Sofia Madsen, Noéline Héritier, Marlen Knobloch

## Abstract

Lipid droplets (LDs) are cellular stores for lipids. These organelles have recently gained interest in neuroscience because they accumulate in various cell types in neurodegenerative diseases. However, their role under physiological conditions is still not fully understood. Classical LD staining methods, which use lipophilic dyes like BODIPY 493/503 (BD493) or antibodies against LD coat proteins, show very few LDs in healthy brain tissue. Our recently developed novel endogenous LD reporter mouse challenges this view. We have been able to detect numerous LDs in healthy brain tissue from both adult and developing mice without staining. To understand why classical staining and endogenous labeling yield different results, we thoroughly investigated the effects of tissue preparation and detergent used in LD detection. We found that BD493 works poorly in brain tissue, while other lipophilic dyes visualize many LDs. We also found that antibody-based LD detection depends on tissue pretreatment and detergent concentration but can reveal a similar number of LDs as observed with the endogenous LD reporter mouse. Taken together, we here present an optimized procedure for LD detection in brain tissue using commercially available dyes and antibodies. Using these methods, we demonstrate that LDs are numerous in healthy brain tissue and substantially accumulate in aged brains in various cell types, including neurons.

## Introduction

Lipid droplets (LDs) are the lipid storage organelles of cells ^1,2^. They contain neutral lipids in the form of triacylglycerides (TAGs) and cholesterol esters (CEs), which are surrounded by a phospholipid monolayer. LDs also have various LD coat proteins, such as members of the perilipin family ^1–3^. Among cell types, adipocytes and hepatocytes have the highest capacity for lipid storage ^4,5^. Adipocytes, in particular, are specialized cells that can store large amounts of lipids in a single large LD (white adipocytes) or multiple smaller LDs (brown adipocytes) ^4^. While these specialized cell types are key to regulate lipid storage, all cell types can form LDs and store neutral lipids to some extent ^6^. LDs were traditionally seen as inert organelles, but recent research has revealed that they are highly dynamic and regulated, with many functions beyond just lipid storage ^6,7^.

LD formation in cells that are not directly involved in lipid metabolism has mainly been associated with disease ^2^. In cases of excess lipid exposure, such as obesity, LDs accumulate ectopically in cells of tissues like the liver, muscle and heart, resulting in gradual dysfunction and disease ^2,8^. Another classic example of excessive LD formation is the transformation of macrophages into foam cells, which are filled with LDs containing CEs. These foam cells eventually contribute to the development of atherosclerosis ^9^.

Recently, there has been increased interest in LDs in the brain, particularly in relation to neurodegenerative disease ^10–12^. Various cell types in the brain, including microglia and astrocytes, have been found to accumulate LDs, particularly in aged mice and in the context of Alzheimer’s disease ^13–17^. These LDs appear to play a role in the pathological processes associated with these conditions. Recent studies have also shown that astrocytes can take in excess lipids from stressed neurons, store them in LDs, and break them down through fatty acid beta-oxidation ^18^. This process may serve as a physiological mechanism for detoxification, and its functioning might be impaired in disease contexts ^18,19^. These findings have triggered a considerable interest in understanding the role of LDs in both normal brain function and pathology ^10–12^. However, traditional staining techniques using lipophilic dyes or antibodies against LD coat proteins have only revealed a small number of LDs in healthy brain tissue, aside from the well-known LD-containing ependymal cells ^20,21^. This raises the question of whether LDs are truly relevant in normal brain physiology or if there are challenges in detecting them in healthy brain tissue.

We have recently developed a novel LD-reporter mouse by tagging the endogenous LD protein perilipin 2 (Plin2) with tdTomato ^22^. This led to fluorescently labelled PLIN2-positive LDs in all tissues expressing Plin2. Surprisingly, our tdTom-Plin2 mouse showed an abundance of LDs throughout the brain in various cell types in healthy 2-month-old mice ^22^. LDs were also numerous and highly dynamic in the developing mouse brain. These findings suggest that the issue lies in revealing LDs in the brain rather than their absence under physiological conditions.

We have therefore revisited classical LD staining approaches, which are effective in other tissues, and applied them to brain tissue from wildtype mice. We have evaluated the impact of brain tissue freezing on LD detection, have compared various commercially available lipophilic dyes, and have assessed the influence of tissue permeabilization when utilizing an antibody against PLIN2.

We here present our optimized methods for visualizing LDs in healthy mouse brain tissue and provide confirmation of their abundance in the mouse brain. We furthermore demonstrate that there is a substantial accumulation of LDs in aged brains, which goes beyond the previously reported accumulation and includes substantial LD deposition in neurons.

## Results

### BD493 works in cells, but does not reveal LDs in young adult mouse brain tissue

We used brains from 2-month-old tdTom-Plin2 mice (Fig. 1A), to either make brain sections for microscopy or extract and culture primary neural stem/progenitor cells (NSPCs). As previously reported, the endogenous tdTom-Plin2 reporter mouse reveals many LDs in the brain of healthy 2-month-old mice ^22^. This is shown here with a representative confocal image of the cortex (Fig. 1B) and the subventricular zone (SVZ) (Suppl. Fig. 1A). However, when using the widely used lipophilic dye BODIPY 493/503 (BD493) to detect LDs, we only found a few LDs in the SVZ and almost no LDs in the cortex of tdTom-Plin2 mice (Fig. 1B, and Suppl. Fig. 1A). This raises questions about whether all the fluorescent tdTomato-positive structures are indeed LDs. Recently, we have shown that primary NSPCs cultured *in vitro* contain a large number of PLIN2-positive LDs ^23^, and that cultured primary NSPCs from tdTom-Plin2 mice also have tdTomato-positive LD-like structures ^22^. Staining of tdTom-Plin2 NSPCs with BD493 revealed a very high co-localisation of the two signals (Fig. 1C), with ring-like tdTomato-positive structures containing BD493 signal. Quantification showed that around 80% of the tdTomato-positive LDs were also positive for BD493 (Fig. 1F), indicating that at least *in vitro*, both the tdTomato and BD493 signal reveal LDs. Interestingly, we consistently detected more LDs with the tdTomato signal than with the BD493 signal, especially smaller structures. This suggests that BD493 might require a certain quantity of neutral lipids to label LDs, whereas the LD-coat protein PLIN2 is able to reveal smaller LDs as well.

**Figure 1:**
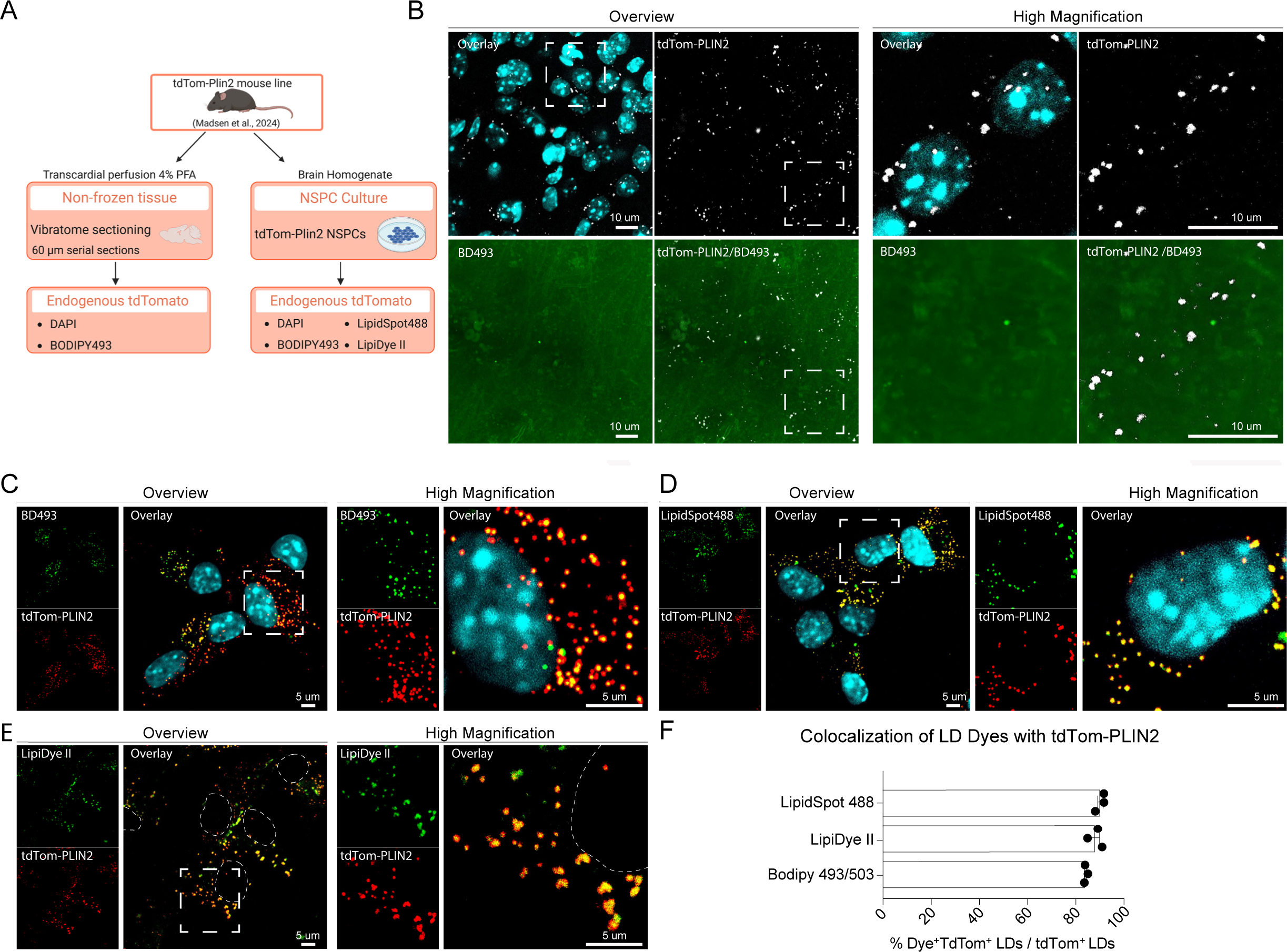
BD493 works in cells but does not reveal LDs in healthy young adult mouse brain tissue. **A)** Scheme illustrating the processing of the brain tissue and staining procedure used for vibratome-derived sagittal brain sections and for NSPCs from tdTom-Plin2 mice. **B)** Representative overview and high magnification confocal images (maximum projections) showing tdTomato (tdTom-PLIN2, in white), DAPI (cyan) and BODIPY493/503 (BD493, in green) in the cortex of 8-week-old tdTom-Plin2 mice. **C, D and E)** Representative overview and high magnification confocal images (maximum projections) showing different lipid dye staining in tdTom-Plin2-derived NSPCs: tdTomato (tdTom-PLIN2, in red), BODIPY493 (BD493, in green), or LipidSpot488 (green), or LipiDye II (green) and DAPI (cyan). Note that due to the excitation of LipiDyeII by 405nm (Suppl. Fig. 1C), nuclei were not counterstained by DAPI in E, but are instead outlined with a dotted line. **F)** Quantification of colocalization puncta of lipid dyes with tdTomato-PLIN2 in NSPCs derived from tdTom-Plin2 mice. The number of LDs is expressed as a percentage of colocalization puncta between LD dye-positive (LD Dye+) and tdTom-PLIN2 positive (tdTom+) LDs per cell over all tdTom+ LDs. Each dot represents data from an individual experiment, with n=3 experiments per group. The data represent the mean value +/- SEM.

### Alternative lipophilic dyes detect a large number of LDs *in vitro* and in brain tissue

While BD493 is one of the most commonly used dye to reveal LDs, there are other lipophilic dyes commercially available that have different chemical properties. We thus wanted to compare how these dyes perform in detecting LDs compared to BD493. We chose LipiDye II, a dye in the green emission spectrum that is designed for live imaging but also works on fixed cells, and a dye class called LipidSpot, which is available in green and far-red (Suppl. Fig. 1B and S1C, of note, far-red is not compatible with tdTomato). These dyes consistently revealed LDs in tdTom-Plin2 NSPCs (Fig. 1D and 1E) and showed a similar or slightly higher percentage of co-localization with the tdTomato signal compared to BD493 (Fig. 1F), suggesting that they are valid alternatives for LD detection.

Next, we assessed and quantified the number of LDs detectable with the different lipophilic dyes in brains from 8-week-old WT mice. After transcardial perfusion with 4% paraformaldehyde (PFA), we used two commonly used tissue processing approaches: vibratome sectioning, which does not require tissue freezing, and microtome sectioning, which is done on frozen tissue that has been incubated in a sucrose solution for better cryopreservation (Fig. 2A). To compare the influence of the tissue sectioning method, we cut the brains in half and processed each hemisphere in parallel (Fig. 2A). Similar to what we had observed with the tdTom-Plin2 mouse brain tissue (Fig. 1C, and Suppl. Fig. 1B), BD493 barely revealed LDs in brain sections from WT mice (Fig. 2B). In contrast, LipiDye II and LipidSpot610 revealed a large number of LDs (Fig. 2C and 2D), very similar to what we had observed with the tdTom-Plin2 mouse. Quantification confirmed that LipiDye II and LipidSpot610 were far superior to BD493 in detecting LDs (Fig. 2E-G). Tissue sectioning did not significantly affect LD detection but freezing of the tissue seemed to slightly reduce the numbers of LDs detected for LipiDye II and BD493, but not for LipidSpot610 (Fig. 2E-G). The background signal was also much lower with LipiDye II and LipidSpot610 compared to BD493 (Suppl Fig. 2).

**Figure 2:**
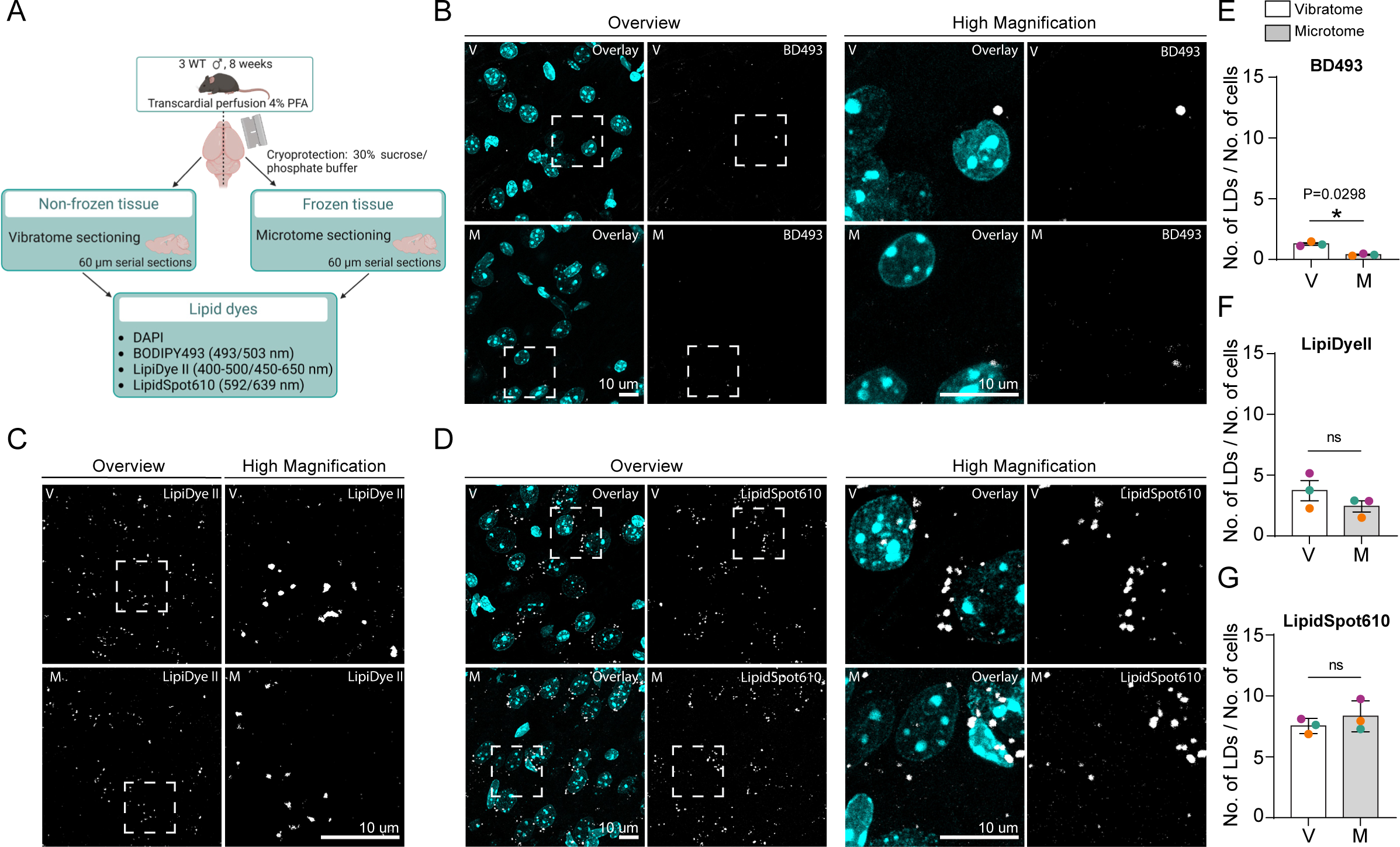
Alternative lipophilic dyes detect a large number of LDs *in vitro* and in brain tissue. **A)** Scheme illustrating the staining procedure in sagittal brain sections of C57BL/6 (WT) mice using both microtome and vibratome sectioning techniques. **B, C and D**) Representative overview and high magnification confocal images (maximum projections) showing different lipid dye staining in microtome (M) and vibratome (V) derived cortical sections of WT mice: BD493 (in white) or LipiDye II (white) or LipidSpot610 (white) and DAPI (cyan). **E, F and G)** The graphs show the quantification of number of LDs detected with BD493, LipiDye II and LipidSpot610, normalized to the number of cells in microtome (M) and vibratome (V)-derived cortical sections of WT mice. Each dot represents data from an individual mouse (color-coded), with n=3 mice per group. The data present the mean value +/- SEM. Paired t-test, p-value: **<0.01, ns = non-significant.

Taken together, these results show that the commonly used lipophilic dye BD493 does not work well for detecting LDs in brain tissue. This might explain why LDs have not been observed in healthy mouse brains by many researchers using BD493. The two alternative dyes perform much better and reveal a large number of LDs. It is worth noting that we are able to successfully stain LDs with BD493 in liver sections using the same staining protocol ^22^, indicating that the issue with BD493 is likely due to specific properties of brain sections.

### Staining outcome using a PLIN2 antibody in adult mouse brain tissue depends on tissue treatment

The tdTom-Plin2 knock-in mouse detects LDs because of the fluorescent tagging of Plin2, which leads to the expression of tdTom-PLIN2 protein ^22^. This mouse is a useful tool for studying LDs without the need for staining in both fixed and live tissues and cells. However, it would also be beneficial to be able to visualize LDs using classical immunohistochemistry approaches in brain tissue. Therefore, we optimized the staining parameters by experimenting with different detergents, detergent concentrations, incubation time, and the addition of detergent to primary and secondary antibody solutions. In our experiments, triton worked best as a detergent when it was only added to the blocking solution and during primary antibody incubation. We compared both vibratome and microtome sections in parallel, using two triton concentrations, 0.3% and 0.15% (Fig. 3A). Overall, we were able to detect a large number of LDs in healthy WT brain sections in all conditions, both in the cortex (Fig. 3B and 3C) and in the SVZ, with clear ring-like structures of large LDs in the SVZ (Suppl. Fig. 3A and 3B). Quantification revealed that the highest number of LDs was detected in vibratome sections with 0.3% triton (Fig. 3D). Reducing the detergent concentration to 0.15% also reduced the number of LDs detected (Fig. 3D). Additionally, the number of LDs detected was lower in microtome sections regardless of the detergent concentration used (Fig. 3D). These findings suggest that treatment of the tissue and the concentration of detergent used can influence the number of LDs that can be detected using immunohistochemistry against PLIN2.

**Figure 3:**
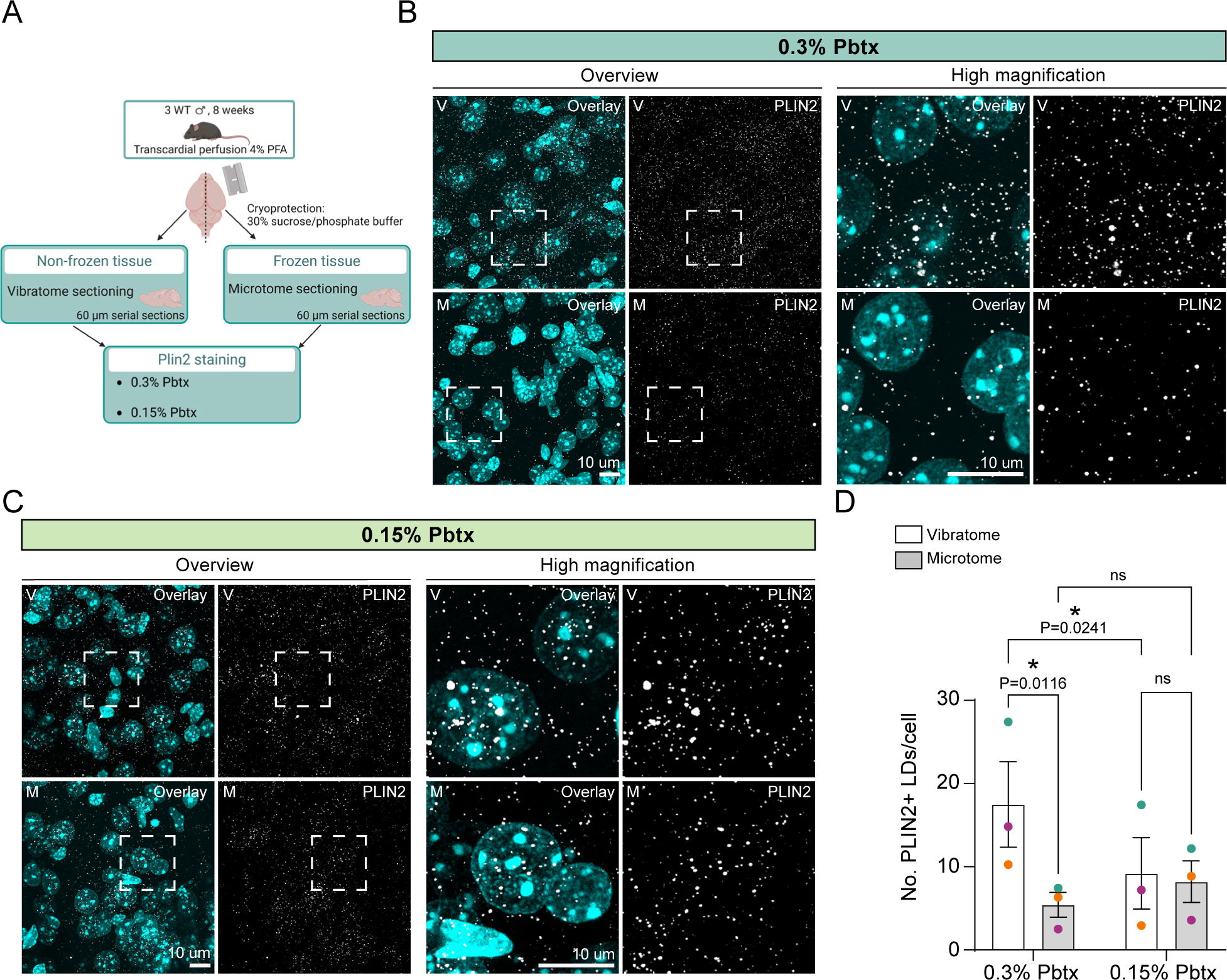
Staining outcome using a PLIN2 antibody in young adult mouse brain tissue depends on tissue treatment. **A)** Scheme illustrating the PLIN2 immunostaining procedures with two different concentrations of phosphate buffer and triton (0.15% and 0.3% Pbtx) in sagittal brain sections of C57BL/6 (WT) mice, using both microtome and vibratome sectioning techniques. **B and C)** Representative overview and high magnification confocal images (maximum projections) showing PLIN2 (white) and DAPI (cyan) immunostaining with 0.3% Pbtx or 0.15% Pbtx in microtome (M) and vibratome (V)-derived cortical sections of WT mice. **D)** Quantification of the number of PLIN2-positive LDs normalized to the number of cells (DAPI), using 0.15% and 0.3% Pbtx in vibratome and microtome sagittal cortical sections of WT mice. Each dot represents data from an individual mouse (color-coded), with n=3 mice per group. The data present the mean value +/- SEM. Two-Way ANOVA followed by Fisher’s LSD test, p-value *<0.05, ns= non-significant.

### Endogenous tdTom-PLIN2 signal does not depend on the tissue sectioning method

To investigate if tissue sectioning affects LD numbers in the endogenous tdTom-Plin2 reporter mouse, we followed the same procedure as with the WT brains (Fig. 2A), using brains from 2-month-old tdTom-Plin2 male mice (Fig. 4A). Surprisingly, freezing the tissue did not impact the number of LDs detected by tdTomato. LD numbers were similar between microtome and vibratome sections, in both the cortex (Fig. 4B and 4C) and the SVZ (Suppl. Fig 4A). These findings suggest that the treatment of the tissue only affects LD detection when combined with the use of detergents, as observed in WT brain sections (Fig. 3D).

**Figure 4:**
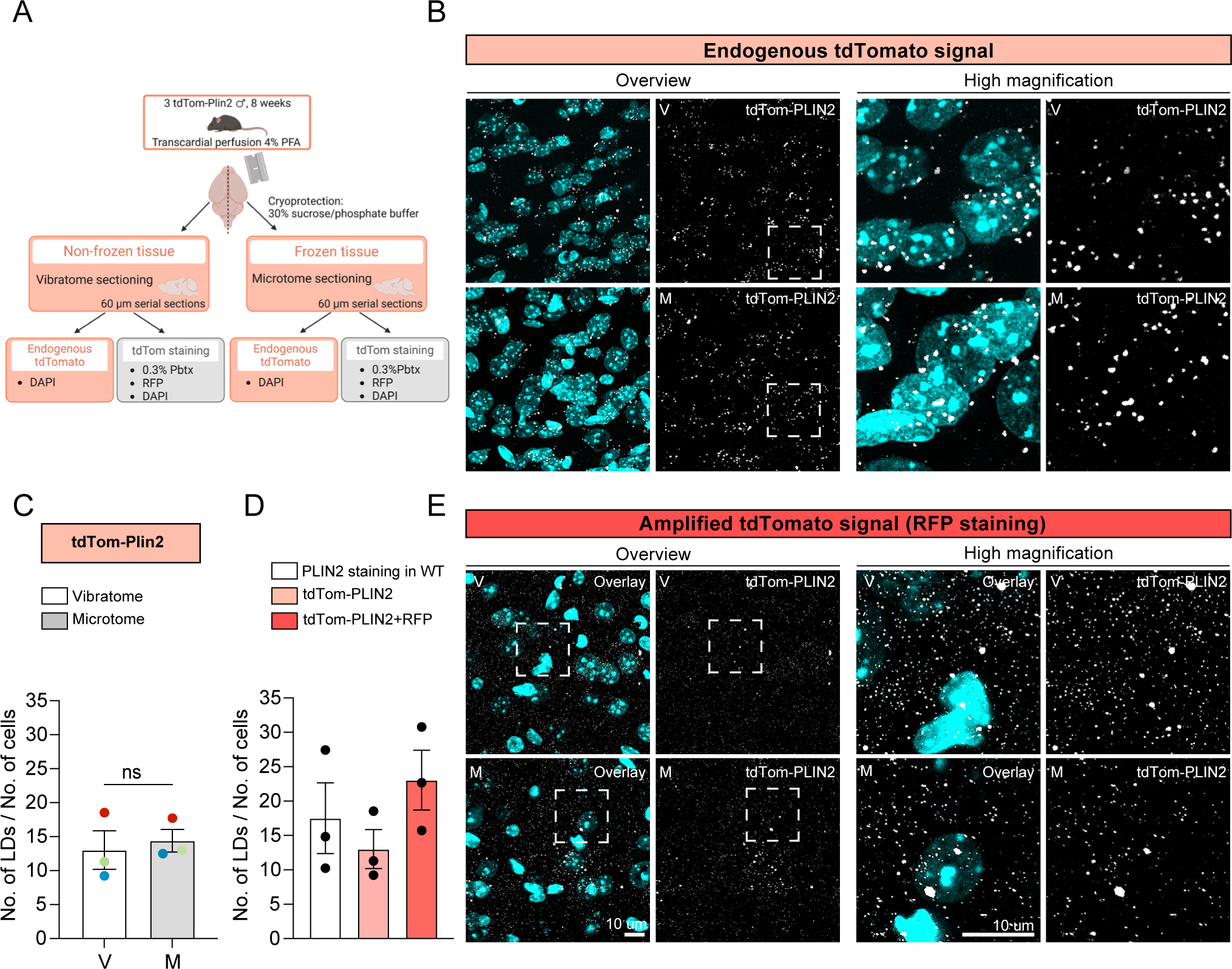
Endogenous tdTom-PLIN2 signal does not depend on the tissue sectioning method. **A)** Scheme illustrating the experimental procedure in sagittal brain sections of tdTom-Plin2 mice, using both microtome and vibratome sectioning techniques. **B)** Representative overview and high magnification confocal images (maximum projections) showing tdTomato (tdTom-PLIN2, white) and DAPI (cyan) in microtome (M) and vibratome (V)-derived cortical sections of tdTom-Plin2 mice. **C)** Quantification of tdTom-Plin2 positive LDs normalized to the number of cells (DAPI) in vibratome and microtome sagittal brain sections of tdTom-Plin2 mice. Each dot represents data from an individual mouse (color-coded), with n=3 mice per group. The data present the mean value +/- SEM. Paired Student t-test, ns=non-significant. **D)** Quantification of the number of LDs in PLIN2-stained, endogenous tdTom-PLIN2 and tdTom-PLIN2 enhanced with RFP-stained LDs, normalized on the number of cells in WT and tdTom-Plin2 mice. Each dot represents data from an individual mouse, with n=3 mice per group. The data represent the mean value +/- SEM. One way ANOVA showed no significance between the groups. **E)** Representative overview and high magnification confocal images depict endogenous tdTom-PLIN2 stained with RFP (white) and DAPI (cyan) in microtome (M) and vibratome (V)-derived cortical sections of tdTom-Plin2 mice.

### Tissue permeabilisation leads to changes in LD size distribution

While the LD numbers in PLIN2-stained WT brains and tdTom-Plin2 brains were comparable (Fig. 4D), we noticed slight differences in the LD pattern in WT brain sections (Fig. 3B and 3C) and tdTom-Plin2 brain sections (Fig. 4B and 4C). Staining for PLIN2 resulted in a higher number of small puncta, and LDs in WT brain sections appeared generally smaller. Therefore, we examined the LD size distribution and confirmed that there was a higher percentage of small LDs in the WT brain sections stained against PLIN2 compared to the tdTom-Plin2 brain sections (Suppl. Fig. 4B). Interestingly, this pattern also emerged when subjecting the tdTom-Plin2 sections to an immunostaining using an antibody against tdTomato (RFP) (Fig. 4D and E, Suppl. Fig. 4C). These results suggest that the immunostaining procedures alters the LD size distribution. The cause of this change, whether it is due to LD shrinkage, lipid leakage during detergent exposure, the inability to detect these smaller LDs without immunostaining amplification, or a combination of these factors, remains to be determined.

### Simultaneous detection of the LD core and LD coat works in tdTom-Plin2 brain sections, but fails with immunostaining of WT brain sections

Stainings of cultured tdTom-Plin2 NSPCs with lipophilic dyes show clear double-labeling, demostrating that the tdTom-Plin2 construct does indeed report LDs, at least *in vitro* (Fig. 1E-G and ^22^). Most of the tdTom-Plin2 signal in brain sections is dot-like, but ring-like structures can also be detected in both tdTom-Plin2 sections (Fig. 4, Suppl. Fig. 4 and ^22^) and PLIN2-stained WT brain sections (Fig. 3, Suppl. Fig.3). To prove that the structures detected in brain sections are indeed *bona fide* LDs, we co-stained tdTom-Plin2 sections with either the lipophilic dye LipidSpot488 or LipiDye II. TdTomato and the lipophilic dyes showed clear co-localization (Fig. 5 A and B), and in larger LDs, a ring-like tdTomato signal surrounded a green lipid dye labelled core (Fig. 5 C and D), suggesting that these are indeed LDs.

**Figure 5:**
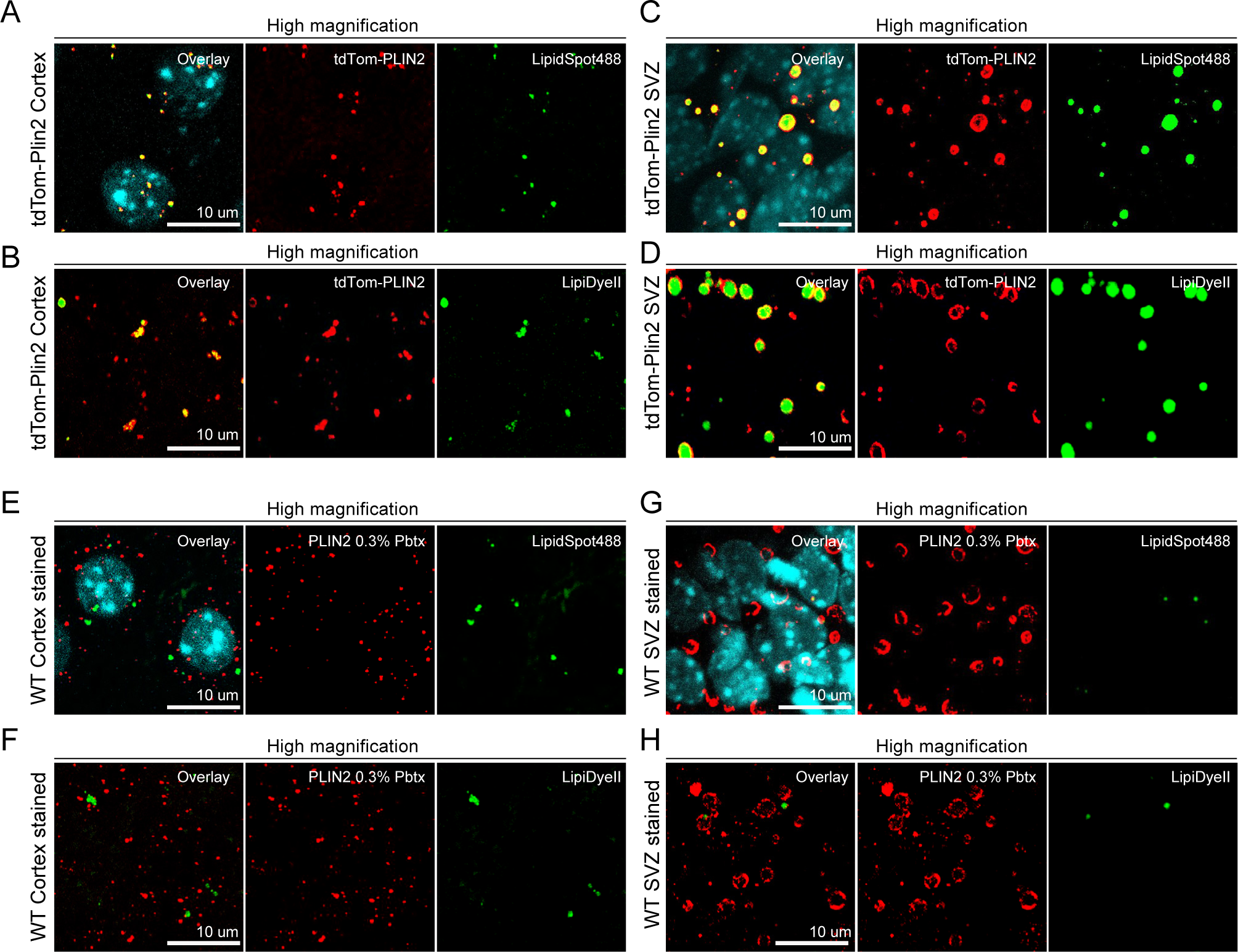
Simultaneous detection of LD core and LD coat works in tdTom-Plin2 brain sections but fails with immunostaining of WT brain sections. **A and B)** High magnification confocal images (maximum projections) showing tdTom-PLIN2 (red), LipidSpot488 or LipiDyeII (green) and DAPI (cyan) staining in vibratome-derived cortical sections of tdTom-Plin2 mice. **C and D)** High magnification confocal images (maximum projections) showing tdTom-PLIN2 (red), LipidSpot488 or LipiDyeII (green) and DAPI (cyan) staining in vibratome-derived sections of the SVZ of tdTom-Plin2 mice. **E and F**) High magnification confocal images (maximum projections) showing PLIN2 (red), LipidSpot488 or LipiDyeII (green) and DAPI (cyan) staining in vibratome-derived cortex sections of WT mice. **G and H)** High magnification confocal images (maximum projections) showing PLIN2 (red), LipidSpot488 or LipiDyeII (green) and DAPI (cyan) staining in vibratome-derived SVZ sections of WT mice.

However, when we combined the LipidSpot488 or LipiDyeII staining with immunohistochemistry against PLIN2 in WT brain sections, the lipophilic dyes performed poorly, despite a clear PLIN2 signal (Fig. 5 E and F). Even in the large ring-like PLIN2 positive LDs in the SVZ, no lipophilic dye signal was detectable (Fig. 5 G and H). This is in strong contrast to the clear signals obtained in WT brain sections when only using the lipophilic dyes without tissue permeabilisation (Fig. 2C and 2D). These data suggest that the lipids in LDs might be washed out during the immunohistochemistry process when using brain tissue, and only the LD coat proteins remain. Alternatively, integration of lipophilic dyes in lipid rich structures might be hindered after the use of detergents. When using the endogenous tdTom-Plin2 LD reporter, there is no need for tissue permeabilisation, and co-labeling is therefore working.

### Substantial accumulation of LDs in brains from 2-year-old tdTom-Plin2- and WT mice

Given the previous literature that LDs accumulate in microglia and astrocytes with aging ^13–17^, we next examined the brains of 2-year-old tdTom-Plin2 mice, focusing on the cortex. To our surprise, almost all cells per field of view had substantial accumulation of LDs. When comparing these aged brains to brains from 2-month-old tdTom-Plin2 mice (P60), the area covered by LDs was almost 9-fold increased (Fig. 6A and 6B). To confirm that this extensive LD accumulation was not only present in the tdTom-Plin2 reporter mouse, but also visible in aged WT mouse brains, we used the optimized staining protocol with an antibody against PLIN2. The cortex of 2-year-old WT mice showed a similar extensive accumulation of LDs in almost all cells per field of view (Suppl. Fig. 5A and 5B). Quantification showed a 7 to 8-fold increase in LDs in the aged brains compared to WT brains at P60, depending on the percentage of detergent used (Suppl. Fig. 5C and 5D). These findings suggest that LD accumulation is substantially increased with age to a much higher extend than previously reported.

**Figure 6:**
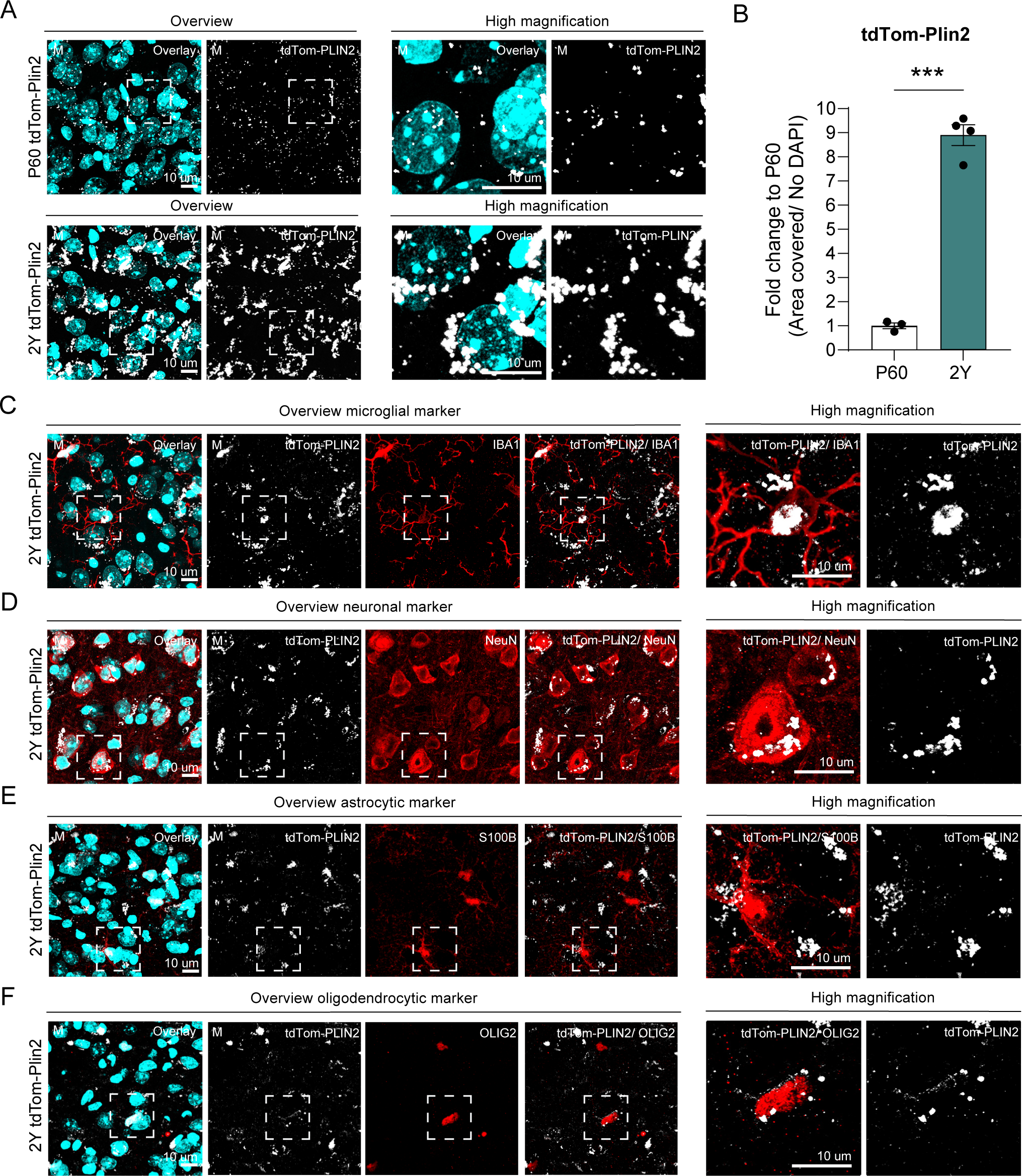
Substantial accumulation of LDs in brains from 2-year-old tdTom-Plin2 and WT mice. **A)** Representative overview and high magnification confocal images (maximum projections) showing tdTomato (tdTom-PLIN2, white) and DAPI (cyan) in microtome sections of the cortex of tdTom-Plin2 mice at P60 (upper panel) and at 2-year of age (lower panel). **B)** Quantification of the area covered by tdTom-PLIN2 positive LDs normalized to the number of cells. Shown is the fold change to P60. Each dot represents an individual mouse, with n=3 mice per group. The data present the mean value +/- SEM. Unpaired Student t-test. p-value ***<0.001. **C to F**) Representative overview and high magnification confocal images (maximum projections) of cortical sections of 2-year-old tdTom-Plin2 mice stained against different cell type markers. Shown is PLIN2 (white) and DAPI (cyan) with the microglial marker IBA1 (**C**, red), the neuronal marker NeuN (**D**, red), the astrocytic marker S100B (**E**, red) and the oligodendrocytic marker OLIG2 (**F**, red).

We next tested whether these LD accumulation in old brains of tdTom-Plin2 and WT mice were also visualized using lipophilic dyes. Interestingly, BD493, which performed poorly in young brain tissue, was able to reveal some of the LDs in the aged mice, however to a much lesser extent than what was seen with the tdTom-PLIN2 signal (Suppl. Fig. 6A and 6B). In contrast, LipiDye II and LipidSpot488 were able to reveal the massive LD accumulation in both WT and tdTom-Plin2 mice and showed very good co-localization with the tdTomato signal (Suppl. Fig. 6C-F).

To address which cell types accumulated LDs, we next co-stained the brain sections from 2-year-old tdTom-Plin2 mice with markers for different cell types, such as microglia (IBA1), neurons (NeuN), astrocytes (S100B) and oligodendrocytes (OLIG2). These stainings revealed LDs in all these cell types, with an especially pronounced accumulation in microglia and neurons (Fig. 6C-F).

## Discussion

LDs in the brain have gained significant interest in the recent years, as they appear to be involved in normal brain function and may also be directly implicated in several neurological diseases ^2,10–12^. However, due to their unique structure, consisting of a phospholipid monolayer and a lipid-rich core, LDs are fragile and tissue processing and detergents used for immunohistochemistry are likely to influence their detection ^24,25^. Therefore, caution must be exercised when selecting a method to visualize LDs, particularly when making statements about their absence, as this could be due to the chosen detection method. Several established techniques exist for detecting LDs, including the use of various lipophilic dyes ^26^ and immunohistochemistry targeting LD coat proteins ^27^. Additionally, several advanced microscopy techniques, that exploit the specific properties of lipids, such as high optical diffraction or specific vibrational characteristics, have been developed for label-free LD detection ^28–30^. However, these microscopy techniques require specialized equipment, making them less accessible. While staining methods work well in cultured cells, detecting LDs in tissues with these methods is more challenging. The brain is a highly lipid-rich tissue, with lipids accounting for over 50% of its dry weight ^31,32^ and the detection of LDs with lipophilic dyes appears to differ from other tissues. Here we demonstrate that the commonly used dye BD493, which performs effectively in cells and various tissues like the liver, does not yield satisfactory results in brain sections (Figure 1). It exhibits a low signal to noise ratio and significant background signal. Surprisingly, other lipophilic dyes with distinct chemical structures, such as LipiDye II and LipidSpot488 and LipidSpot610 perform much better in brain sections compared to BD493 (Figure 2). We confirmed that all dyes performed similarly well in cultured NSPCs *in vitro*, ruling out a general issue with our BD493 protocol (Figure 1). The reason for this discrepancy in dye performance is unclear, but it underscores the significant impact that the choice of lipophilic dye can have on results. Tissue penetrance and binding properties may vary among the dyes. Given the lipid-rich nature of brain tissue, certain lipophilic dyes may have a greater tendency to interact with myelin, potentially reducing their accumulation in LDs. Indeed, we observed clear fiber tracks when using BD493, reminiscent of myelinated axons. Interestingly, recent studies have revealed that the presence of crystalline or liquid cores in LDs depends on the ratio of TAGs to CEs ^33^. Whether this phenomenon influences LD detection in the brain and whether different lipophilic dyes perform differently depending on the core structure remains to be determined.

As LDs are rather fragile organelles, tissue treatment is likely to influence their detection. We maintained the initial fixation of the tissue using 4% PFA. However, it remains to be investigated whether different tissue fixation methods will also affect LD detection. In this study, we evaluated two parameters of tissue treatment, freezing or not freezing the tissue for sectioning, and the percentage of triton used as a detergent during immunohistochemistry (Figure 2). To prevent water crystal formation during the freezing process, we utilized the well-established cryopreservation strategy of immersing the brain in 30% sucrose solution before freezing. The tdTom-Plin2 mouse ^22^ allowed us to distinguish the effects of tissue treatment and detergent concentration, as the endogenous fluorescent LDs are detected without the need for immunohistochemistry. Interestingly, freezing or not freezing the tissue prior to cutting only influenced LD detection when immunohistochemistry was performed (Figure 3), but no significant difference was observed in the tdTom-Plin2 sections (Figure 4). This suggests that LDs are not inherently destroyed by tissue freezing processes, but the subsequent use of detergent can affect LD detection. Lipids, which are notably not crosslinked by PFA, thus not fixed, may leak out and alter LD detection. Our findings support this leakage hypothesis, as the LD coat proteins were more consistently detected than the LD core (Figure 5). Additionally, the simultaneous detection of LD coat and LD core was only achievable with the tdTom-Plin2 reporter mouse, where no detergents were required to reveal the LD coat protein. However, in WT mice, simultaneous detection of LD coat and LD core was not possible, likely due to lipid leakage after the immunochemistry procedure (Figure 5 and suppl. Figure 5). In line with this, we also observed less reliable lipophilic dye stainings with larger LDs, such as those found in the SVZ, suggesting that lipids from larger LDs may be more prone to leakage. We also noted that when using lipophilic dyes, it is important to proceed to the microscope within one week, as the dyes can leak even when the tissue or cells are mounted on glass carriers and embedded with antifade solution.

While our data clearly demonstrate that LDs are also abundant in the healthy brain and at young age, the specific role they play is largely unknown. Using the tdTom-Plin2 LD reporter mouse, we have shown that various types of brain cells contain LDs, in varying numbers and sizes ^22^. These findings suggest that having a certain number of LDs seems to be physiologically normal. Indeed, the recent discovery that astrocytes take up excessive lipids from neurons and store them in LDs ^18^ highlights the close metabolic interplay between these two cell types. However, we are only beginning to understand the physiological function of LDs in the brain. Therefore, further research is necessary to uncover their dynamics and to understand their importance for normal brain function.

Our protocol also demonstrates a substantial accumulation of LDs with aging. LD accumulation with aging has been previously reported ^13–17^. However, comparing the results obtained in this study with the previously published studies shows that LD accumulation might have been clearly underestimated (Figure 6). Since we only compared P60 and 2-year-old mice, it is still unclear whether accumulation occurs gradually over time or if there is a specific age at which this increase becomes visible. The global accumulation of LDs and its magnitude is surprising, as it appears to impact most cells in the cortex, including neurons. Further studies are required to better characterize which cells accumulate LDs, why they accumulate them and how this accumulation affects their functionality. Furthermore, it will be interesting to study whether there are regional and temporal differences in the accumulation of LDs with age.

Taken together, our data show that LDs are abundant in the healthy adult mouse brain and accumulate substantially with aging. LDs can be detected using our novel endogenous tdTom-Plin2 reporter line ^22^, as well as lipophilic dyes or immunohistochemistry. We show that tissue treatment, selection of the dyes and detergent concentration all have a clear influence on the detection of LDs. Therefore, care has to be taken and researchers should utilize several methods to assess LDs in the brain. Our results provide a basis for other researchers interested in studying LDs in the brain, helping them choose an appropriate staining method. The massive accumulation of LDs with aging, which we here describe using the tdTom-Plin2 mouse and the optimized staining methods, also raises questions, such as whether and how this might disturb normal brain function and/or influence neurodegenerative disease progression. Having tools at hand to reliably visualize LDs in the brain should facilitate research aiming at solving these important questions.

## Material and Methods

### Animals

All experiments involving animals were conducted in accordance with the Swiss law and received prior approval from the local authorities (Cantonal Veterinary office, Vaud, Switzerland). TdTom-Plin2 mice were generated as described ^22^. 8-week-old and 2-year-old tdTom-Plin2 mice and C57BL/6Rj WT mice (Janvier, France) were used in this study. All mice were kept under standard housing on a 12:12 h light/dark cycle, in ventilated cages with ad libitum food and water. Male mice were used for all experiments, except for NSPC extraction and for the aged 2-year-old tdTom-Plin2 mice, where 3 females and 1 male were used due to the limited availability of old tdTom-Plin2 mice.

### NSPC extraction and expansion

Adult mouse NSPCs were isolated from the SVZ of two 8-week-old tdTom-Plin2 females mice as previously described ^34^. In brief, mice were shortly anaesthetized with isoflurane, followed by decapitation. SVZ were micro-dissected, and a single cell suspension was generated using the papain-based MACS Neural Tissue Dissociation Kit (#130-092-628, Milteny) and the GentleMacs Dissociator (Milteny), according to the manufacturer’s instructions. Myelin removal was performed using the MACs myelin removal beads (#130-096-731, Milteny,) and a QuadroMACS Separator (#130-090-976, Milteny) according to the manufacturer’s instructions. The obtained cells were cultured as neurospheres in DMEM/F12/GlutaMAX (#31331-028, Gibco) with B27 (#17504044, Gibco), 20 ng/ml human EGF (#AF-100-15, PeproTech), 20 ng/ml human basic FGF-2 (#100-18B, PeproTech), and 1x PSF (#15240062, Gibco). Medium was changed every 2-3 days. The neurospheres were expanded for 5 passages to remove progenitors and other proliferating cells. After 5 passages, cells were changed to the following culture medium: DMEM/F12/GlutaMAX (#31331-028, Gibco), N2 (#11520536, Gibco), 20 ng/ml human EGF (#AF-100-15, PeproTech), 20 ng/ml human basic FGF-2 (#100-18B, PeproTech), 5mg/ml Heparin (#H3149-50KU, Sigma) and 1x PSF (#15240062, Gibco). All the *in vitro* experiments were done on passages 7-15.

### Cell culture

NSPCs were grown on uncoated plastic cell culture dishes for expansion (#430167, Corning, TC-treated). Cells used for experiments were plated on glass coverslips (#10337423, Fisher) coated with poly-L-ornithine (#P3655, Sigma) and laminin (#L2020-1MG, Sigma). Proliferating NSPCs were kept in DMEM/F12/GlutaMAX (#31331-028, Gibco) complemented with N2 (#11520536, Gibco), 20 ng/ml human EGF (#AF-100-15, PeproTech), 20 ng/ml human basic FGF-2 (#100-18B, PeproTech), 5 mg/ml Heparin (#H3149-50KU, Sigma) and 1X PSF (#15240062, Gibco). Medium was changed every 2-3 days.

### Tissue preparation

#### Perfusion

All experimental mice were deeply anesthetized through intraperitoneal injection (i.p.) of pentobarbital (150mg/kg) and subsequently intracardially perfused first with ice cold 0.9% saline until no blood remained, then perfused with 40 mL fresh 4% paraformaldehyde (PFA, #19208, Electron Microscopy Sciences, EMS) in 0.1M phosphate-buffered saline (pH 7.4). The brains were post-fixed overnight at 4°C in PFA. After post-fixation, the brains were divided in two hemispheres and stored at 4C in PBS supplemented with 0.02% sodium azide (RTC000068, Sigma).

Half of the hemispheres were cut in sagittal sections of 60µm using a vibratome (Leica) and stored at 4°C in 1X PBS supplemented with 0.02% sodium azide (#RTC000068, Sigma Aldrich). The other hemispheres were incubated in sucrose 30% in phosphate 0.1M at 4°C for 48h for cryoprotection. They were then frozen on a dry ice-cooled metal stage and cut in 60um sagittal sections on a sliding microtome (Leica). The sections were stored in cryopreservation solution (25 % ethylene glycol, 25 % glycerol in 0.05 M phosphate buffer) at 4 °C.

### Immunohistochemistry

For immunohistochemical analysis, sections were washed three times in 1X PBS for 5 min on orbital shaker at RT. Sections were permeabilized for 1h in 1X PBS containing 0.3% or 0.15% Triton X-100 (#X100-100ML, Sigma), and 10% donkey serum (#S30-100ml, Merck) and then immunolabeled for 48h at 4°C on an orbital shaker, using the following primary antibodies: Rabbit-PLIN2 (1:600, Abcam, ab52356), rabbit-RFP (1:600, ab62341). After the primary antibody incubation, the sections were washed again three times in 1X PBS for 10 min and incubated for 2h at RT with fluorescent secondary antibodies (1:300, AlexaFluor, anti-rabbit 488, donkey anti-rabbit 594, Jackson, 711-545-152) diluted in 1X PBS. Finally, nuclei were counterstained with 4’, 6-diamidino-2-phenylindole (DAPI) (Invitrogen, 1:10000) for 15 min in 1X PBS, washed two times in 1X PBS. Sections were mounted on 25×75×1mm Superfrost Plus adhesion microscopes slides (#J1800AMNZ, Epredia) using FluorSave^TM^ reagent (#345789, Sigma).

### Lipophilic dye staining in NSPCs and brain tissue

LDs in NSPCs were detected as following: Cells were fixed with 4%PFA (pre-warmed at 37°C) for 20 min at RT, followed by two washes with 1X PBS for 10 min. Subsequently, the fixed cells were stained for 1h at RT with the following lipophilic dyes, diluted in 1X PBS: BODIPY 493/503 4,4-DIFLUORO-1,3,5,7,8-PE (Fisher Scientific 11540326) (1:1000), LipidSpot 488 or 610 (VWR 70065-T or 70069-T) (1:1000), LipiDye II (Anawa, FDV-0027) (1:1000). Cells were subsequently washed three times in 1X PBS for 5 min. After that, nuclei were counterstained with 4’, 6-diamidino-2-phenylindole (DAPI) (Invitrogen, 1:5000), for 5 min in 1X PBS, and then washed twice with 1X PBS for 5 min before mounting with a home-made PVA-DABCO-based mounting medium (Glycerol (Sigma 49782-1L, 24%), PVA (Sigma P8136-250g, 9.6%), 96mMTrisHCl, DABCO (Sigma D27802-100G, 2.5%)).

LDs in brain sections were visualized using the same lipophilic dyes. Briefly, brain sections were initially washed twice for 5 min each in 1X PBS. Subsequently, they were incubated for 2h at RT on an orbital shaker with the different dyes and DAPI simultaneously, using the concentrations indicated above. Following the incubation with the fluorescent dyes and DAPI, the sections were washed twice in 1X PBS for 5 min and were then mounted with a home-made PVA-DABCO-based mounting medium.

### Immunohistochemistry of PLIN2 with LD fluorescent dyes

For immunohistochemical analysis of PLIN2 and fluorescent dyes, sections were washed three times in 1X PBS for 5 min on an orbital shaker at RT. Then, sections were permeabilized for 1h in 1X PBS containing 0.3% or 0.15% Triton X-100 and 10% donkey serum (Merck, #S30-100ml). Thereafter, they were immunolabeled for 48h at 4°C on an orbital shaker, using the following primary antibodies: Rabbit-PLIN2 (1:600, Abcam, ab52356). Following the primary antibody incubation, the sections underwent three additional washes in 1X PBS for 5 min each and were then incubated for 2h at RT with fluorescent secondary antibodies (AlexaFluor, anti-rabbit 488, donkey anti-rabbit 594, Jackson, 711-545-152) diluted in 1X PBS. Finally, nuclei and LDs were counterstained with DAPI and the following fluorescent dyes: BODIPY 493/503 4,4-DIFLUORO-1,3,5,7,8-PE (Fisher Scientific 11540326, dilution 1:1000 in 1X PBS); LipidSpot488 (VWR 70065-T, dilution 1:1000 in 1X PBS); LipidSpot610 (VWR 70069-T, dilution 1:1000 in 1X PBS); LipiDye II 488 (Anawa, FDV-0027, dilution 1:1000 in 1X PBS). After 2h of incubation, the sections were washed twice in 1X PBS and then mounted with a home-made PVA-DABCO-based mounting medium.

### Confocal microscopy acquisition and image analysis

All images were collected on a Leica confocal imaging system (TCS SP8) with 63× (0.75 and 2 NA) oil immersion objective. For the quantification of LDs, serial sections of the cortex and SVZ were used. Briefly, Z-stacks were taken at 0.3 µm intervals, and the number and the diameter of LDs in the P60 mouse brain were measured using IMARIS software. The “Spot” function was used to perform a 3D reconstruction of all LDs, with a minimum volume threshold set at 0.05 µm³.

Three images per sections and several sections per regions were taken from 3 mice per group. For the quantification of colocalization of Lipid Dyes with tdTom-Plin2 in NSPCs, 3 coverslip per condition were stained, 3 images per coverslip were taken using digital zoom 3x and all Lipid Droplets in the field of view were analysed using IMARIS software. Briefly, “Spots” function was created for lipid droplets stained with Lipid Dyes and for tdTom-Plin2 positive lipid droplets. Colocalization between two created spot sets was analysed using “Spot Colocalization” function.

To quantify the LDs in the aged brain, we used the freely-available, interactive machine-learning software called ilastik ^35^ (Version 1.4.0.post1, released on November 10, 2023). This method was chosen because of the clustering of LDs in the two-year-old mice, which did not allow to separate individual LDs. Representative images for each condition were loaded as input images to manually train the software in recognizing LDs from background. After training, simple segmentation images were extracted from ilastik. Then, using the “Analyze particles” tool in Fiji, the percentage of area covered was measured. To account for different number of cells, the percentage of area covered was normalized to the number of nuclei stained with DAPI.

### Statistical Analyses

To compare the data obtained from the two hemispheres undergoing different tissue sectioning, paired t-test were used. For comparing the influence of tissue sectioning and detergent use, a 2-way ANOVA followed by Fishers LSD test was used. For comparing the LD numbers in WT and tdTom-Plin2 unstained and stained tissue, an ordinary 1-way ANOVA was performed.

All analyses were performed using GraphPad Prism 10.1.2 software (GraphPad software).

### Illustration software

For illustration schemes, Biorender software (Biorender, 2021) and Adobe Illustrator (version 25.0, Adobe Inc) was used.

## Author contributions

FP and AR performed experiment, analyzed and visualized the data. DP performed the experiments in NSPCs, SM and MK developed the tdTom-Plin2 mouse model, NH performed experiments and analyzed data. MK and FP developed the concept and wrote the manuscript, with input from all authors. MK provided the financial means to perform this project.

## Acknowledgements

We thank the Cellular Imaging Facility and the animal facility of the University of Lausanne for technical support. This work was supported by funding from the University of Lausanne and the Swiss National Science Foundation (grant # 31003A_175570, M.K).

---

**Supplementary Figure 1:**
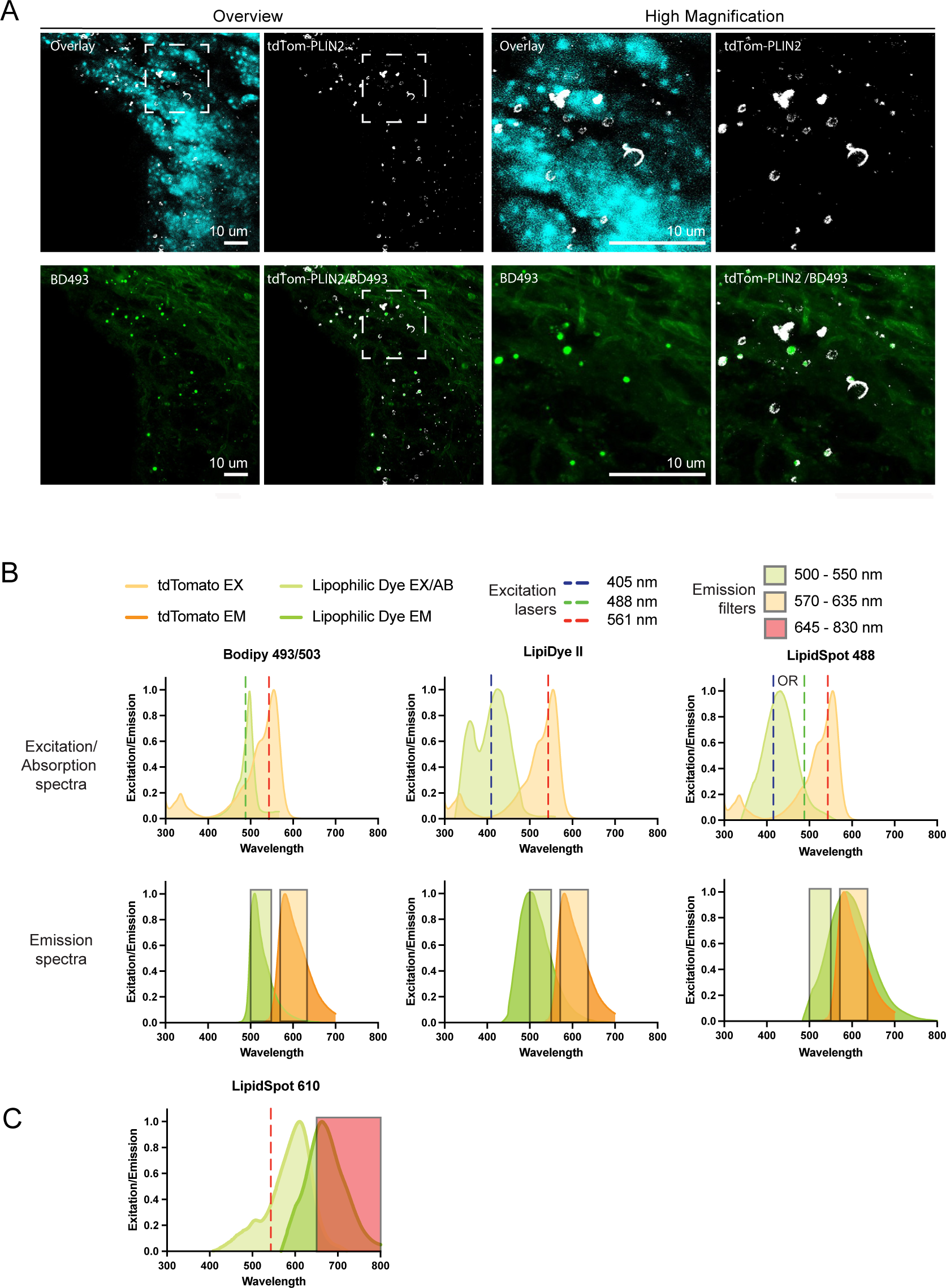
BD493 works in cells but does not reveal LDs in young adult mouse brain tissue, related to Figure 1. **A)** Representative overview and high magnification confocal images (maximum projections) display tdTomato (tdTom-PLIN2, in white), BODIPY 493/503 (BD493, in green) and DAPI (cyan) in the subventricular zone (SVZ) of 8-week-old tdTom-Plin2 mice. **B and C)** Graphs illustrate the excitation and absorption spectra of BODIPY 493/503, LipiDye II, LipidSpot 488 and LipidSpot610. The spectra for tdTomato, BD493/503, LipidSpot 488 and LipidSpot610 were generated using fpbase.org. The spectra for LipiDye II were adapted from data provided by https://www.diagnocine.com/Product/LipiDye-II-Lipid-dye-Droplet-Staining/67496. Note that LipidSpot610 cannot be used together with tdTomato due to the spectral overlap, and that LipiDyeII is also excited by the 405nm laser, thus cannot be combined with DAPI.

**Supplementary Figure 2:**
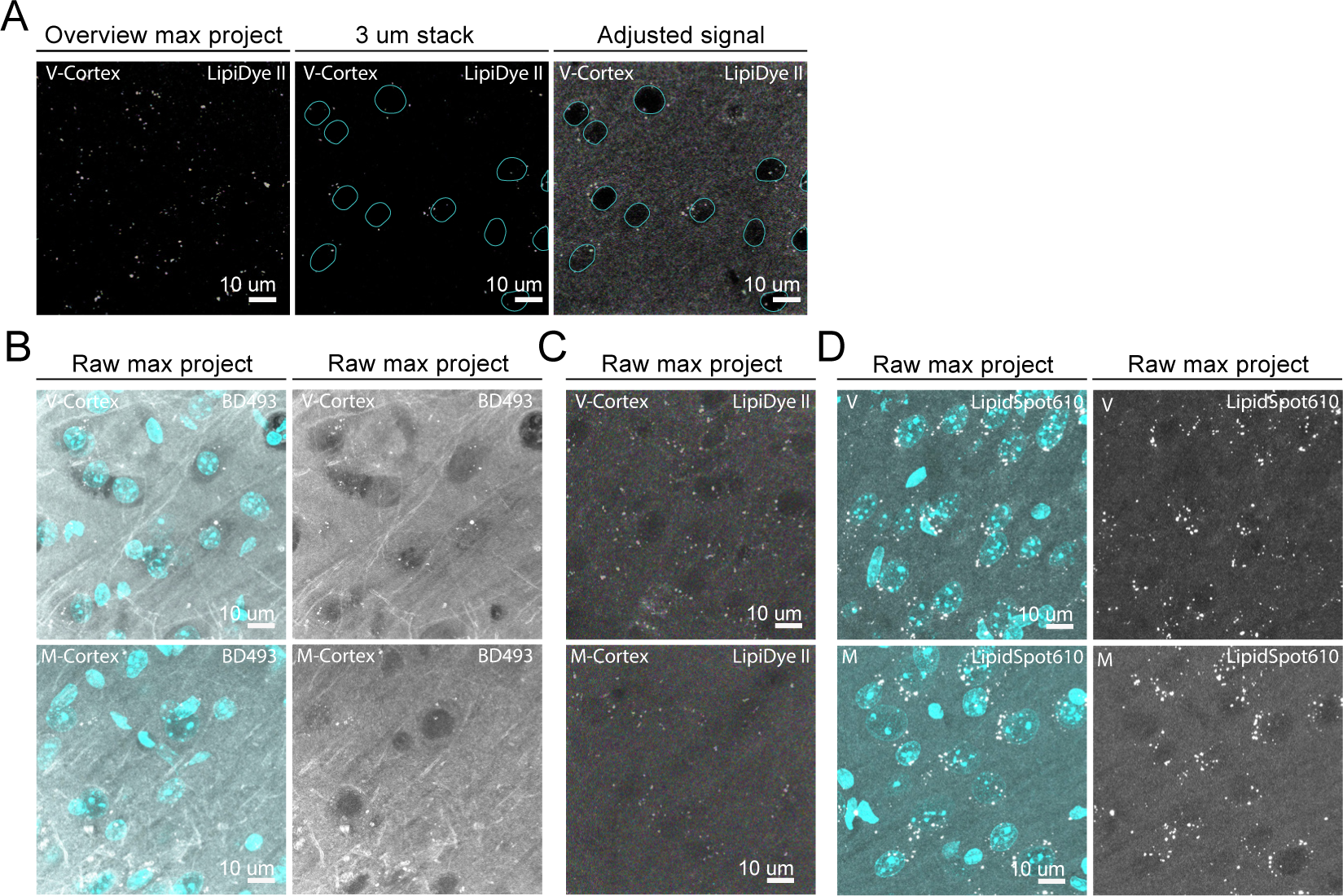
Alternative lipophilic dyes detect a large number of LDs *in vitro* and in brain tissue, related to Figure 2. **A)** Representative confocal images show an overview (maximum projection), a 3 μm projection and the adjusted signal of LipiDye II (white) staining in vibratome sagittal cortical sections of 8-week-old WT mice, to illustrate how the signal can be used to determine cell nuclei. Due to the excitation spectra, DAPI cannot be used. **B, C and D)** Representative confocal images show the raw maximum projections of BD493 (white) or LipiDye II (white) or LipidSpot610 (white) and DAPI (cyan) staining in microtome (M) and vibratome (V)-derived cortical sections of WT mice.

**Supplementary Figure 3:**
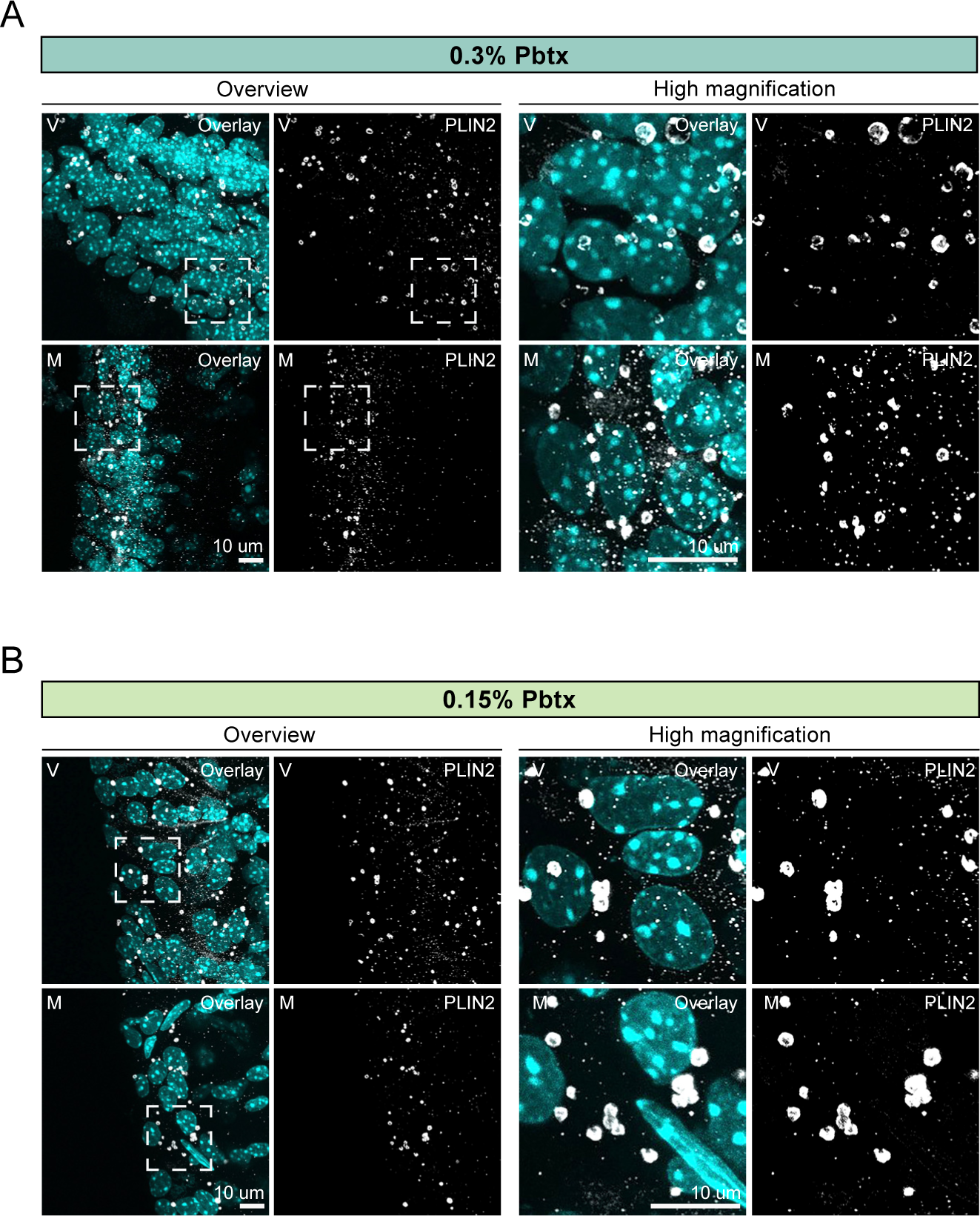
Staining outcome using a PLIN2 antibody in young adult mouse brain tissue depends on tissue treatment, related to Figure 3. **A and B)** Representative overview and high magnification confocal images (maximum projections showing PLIN2 (white) and DAPI (cyan) immunostaining with 0.3% or 0.15% Pbtx in microtome (M) and vibratome (V)-derived SVZ sections of WT mice.

**Supplementary Figure 4:**
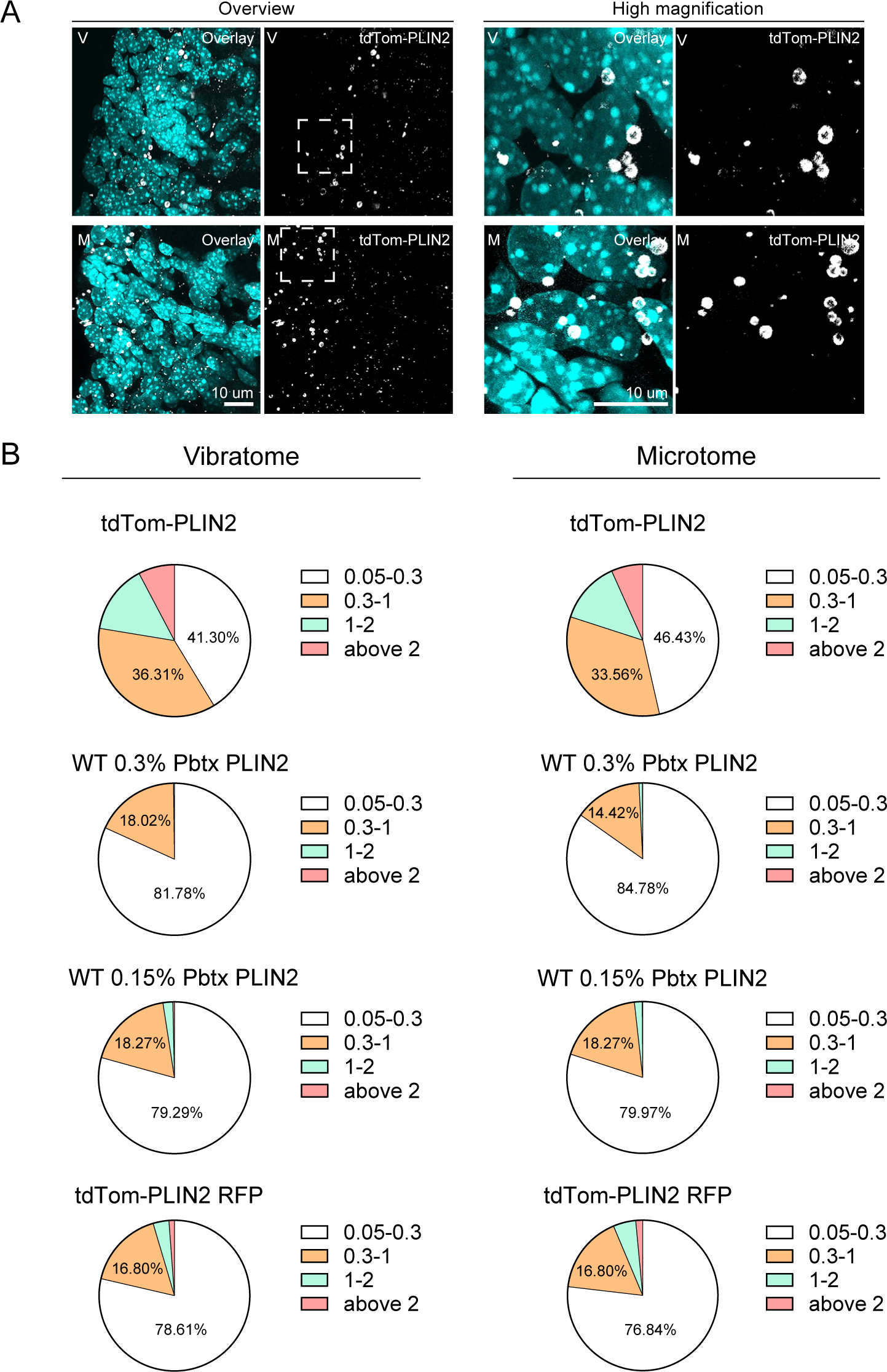
Endogenous tdTom-PLIN2 signal does not depend on the tissue sectioning method, related to Figure 4. **A)** Representative overview and high magnification confocal images (maximum projections) showing tdTomato (tdTom-PLIN2, in white) and DAPI (cyan) staining in microtome (M) and vibratome (V)-derived SVZ sections of tdTom-Plin2 mice. **B)** Pie-charts illustrate the percentages of LD diameters, ranging from 0.05 μm to above 2 μm, across the different experimental conditions.

**Supplementary Figure 5:**
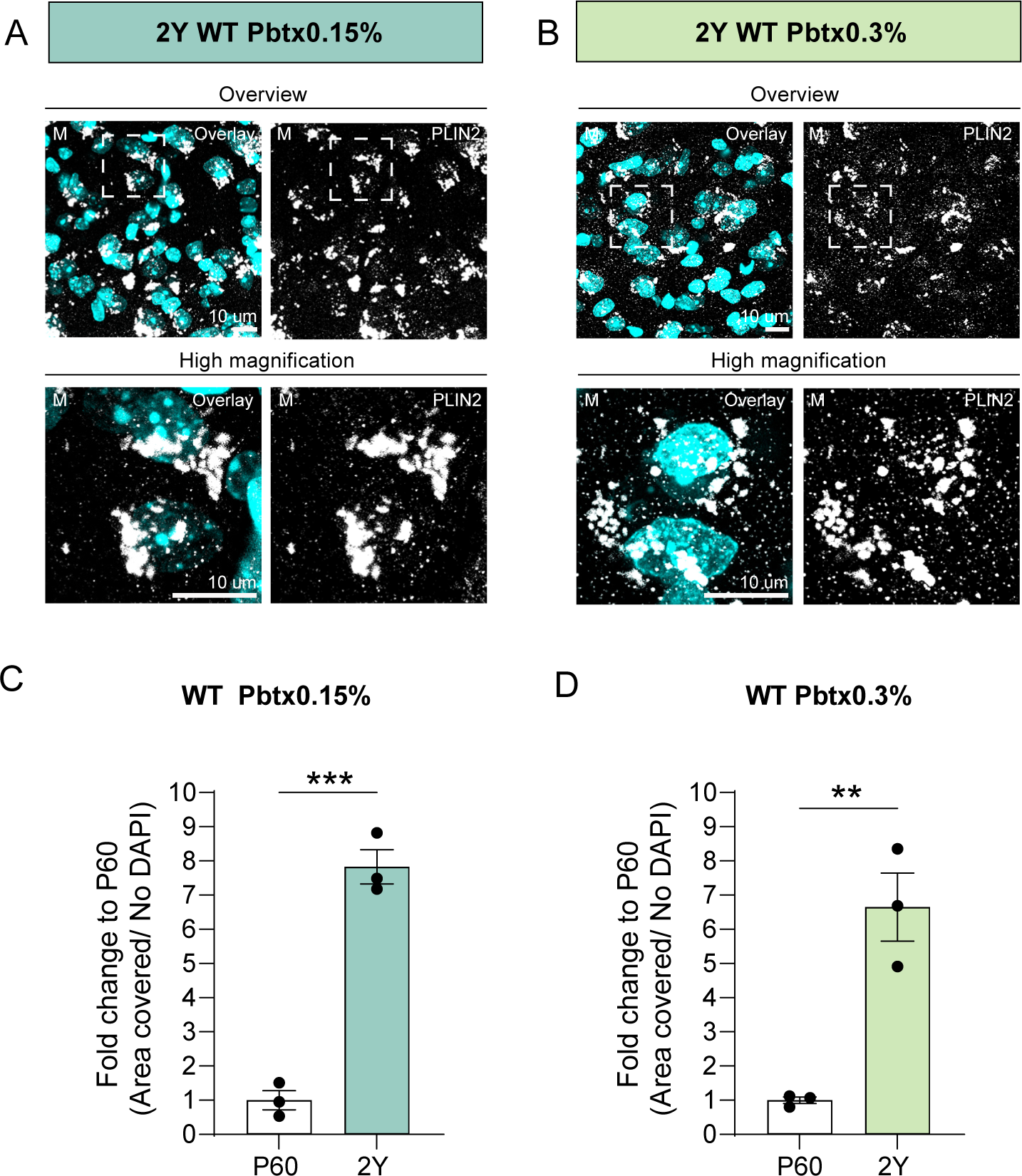
Substantial accumulation of LDs in brains from 2-year-old tdTom-Plin2 and WT mice, related to Figure 6. **A and B)** Representative overview and high magnification confocal images (maximum projections) showing PLIN2 (white) and DAPI (cyan) in microtome sections of the cortex of WT mice at 2-year of age, stained using either 0.15% or 0.3% Pbtx. **C and D)** Quantification of the area covered by PLIN2 positive LDs normalized to the number of cells. Shown is the fold change to P60. Each dot represents an individual mouse, with n=3 mice per group. The data present the mean value +/- SEM. Unpaired Student t-test. p-value ***<0.001, **<0.01.

**Supplementary Figure 6:**
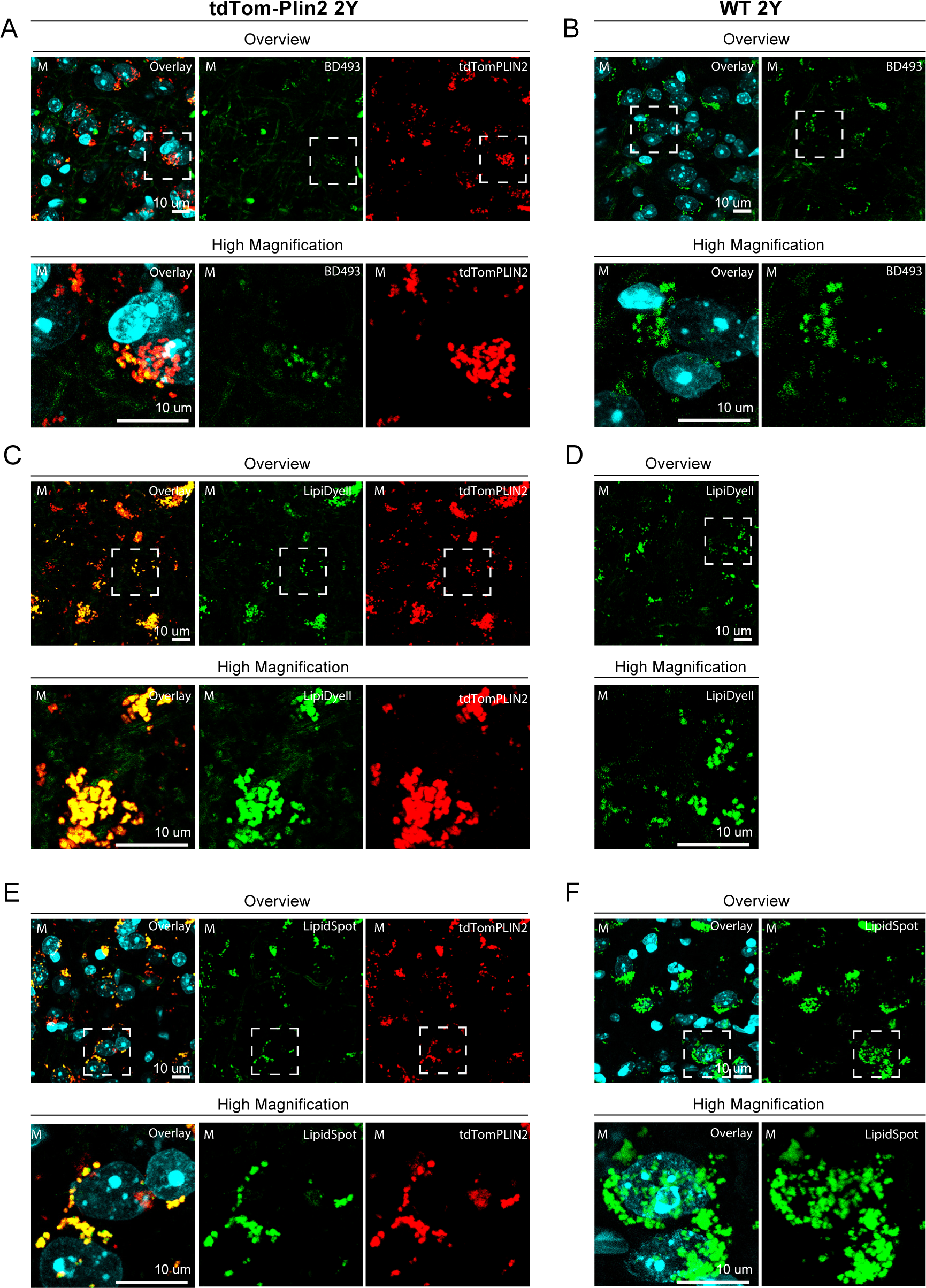
Substantial accumulation of LDs in brains from 2-year-old tdTom-Plin2- and WT mice, related to Figure 6. **A, B and C)** Representative overview and high magnification confocal images (maximum projections) showing the staining with different lipophilic dyes in cortical sections of 2-year-old tdTom-Plin2 mice. Shown is tdTomato (tdTom-PLIN2, red), DAPI (cyan), BD493 (**A**, green), LipidDye II (**B**, green) and LipidSpot488 (**C**, green). **D, E and F)** Representative overview and high magnification confocal images (maximum projections) showing the staining with different lipophilic dyes in cortical sections of 2-year-old WT mice. Shown is DAPI (cyan), BD493 (**D**, green), LipidDye II (**E**, green) and LipidSpot488 (**F**, green). Note that BD493 detects much less LDs than the other two lipophilic dyes.

